# SIK2: A novel negative feedback regulator of FGF signaling

**DOI:** 10.1101/2023.12.26.573381

**Authors:** Gamze Kuser-Abali, Asli Ugurlu-Bayarslan, Yeliz Yilmaz, Ferruh Ozcan, Funda Karaer, Kuyas Bugra

**Affiliations:** Department of Molecular Biology and Genetics, Bogazici University, 34342 Bebek, Istanbul, Turkey; Faculty of Medicine Nursing & Health Sciences, The Central Clinical School, Monash University, Melbourne, Victoria, Australia; Department of Biology, Kastamonu University, 37150 Kastamonu, Turkey; Department of Medical Biology, Dokuz Eylul University Medical School, Inciralti, 35340 Izmir, Turkey; Department of Molecular Biology and Genetics, Gebze Technical University, 41400 Gebze, Kocaeli, Turkey; Ministry of Education, Turkey; Life Sciences Center, Bogazici University, 34342 Bebek, Istanbul, Turkey

**Keywords:** SIK2, FGF2 signaling, Ras/ERK pathway, control of proliferation

## Abstract

A wide range of cells respond to FGF2 by proliferation via activation of the Ras/ERK pathway. In this study we explored the potential involvement of serine/threonine kinase SIK2 in this cascade within retinal Müller glia. We found that SIK2 phosphorylation status and activity is modulated in an FGF2-dependent manner, possibly via ERK. With SIK2 downregulation we observed enhanced ERK activation with delayed attenuation and increased cell proliferation, while SIK2 overexpression hampered FGF-dependent ERK activation. In vitro kinase and site directed mutagenesis studies indicated that SIK2 targets the pathway element Gab1 on Ser266. This phosphorylation event weakens Gab1 interactions with its partners Grb2 and Shp2. Collectively, our results suggest that during FGF dependent proliferation process ERK-mediated activation of SIK2 targets Gab1, resulting in downregulation of the Ras/ERK cascade in a feedback loop.

## Introduction

Fibroblast Growth Factors (FGFs), a family of 23 members in mammals, are critical in regulating a wide spectrum of biological processes including cell proliferation, differentiation, migration and survival [20,53]. FGFs elicit cellular responses by interacting with FGF receptors (FGFRs) that are members of the receptor tyrosine kinase (RTK) superfamily [20,42]. FGF/FGFR signaling is highly regulated by a multitude of feedback mechanisms [66], and its dysregulation has been linked to a wide range of pathological events including developmental disorders and tumorigenesis [6].

The two most prominent pathways activated by FGFs are the Ras/ERK and PI3K/Akt axes [20,48]. Activation of both of these pathways is initiated by phosphorylation of FGF Receptor Substrate 2 (FRS2) on specific tyrosine residues by FGFR, facilitating the formation of a FRS2/Grb2/Gab1 ternary complex proximal to the receptor. Tyrosine phosphorylated Grb2 is instrumental in the recruitment of Sos to the complex, promoting the Ras/ERK cascade [55]. Since FGFR has no docking site for PI3K, receptor-proximal recruitment of the p85 subunit of PI3K by tyrosine phosphorylated Gab1 is crucial for the activation of the AKT signaling [52]. In regards to function, proliferation is predominantly associated with Ras/ERK activation and survival with PI3K/AKT axis in diverse cell types including retinal Müller glia [28,35,36,52].

The docking protein Gab1, through its interaction with Shp2, also contributes to sustained and/or enhanced ERK activation in response to multiple growth factors [32,69]. Shp2 has been proposed to dephosphorylate Gab1 on certain tyrosine residues thus interfering with the recruitment of p120RasGAP, a negative regulator of Ras activation [3,18]. In chondrocytes inhibition of the Shp2/Gab1 association decreased FGF2-induced ERK activation [37]. In a recent study, Shp2 was proposed to mediate a basal level of Ras/ERK signaling in Müller glia [13]. Though there exist a large number of potential serine/threonine phosphorylation sites on Gab1, information on their physiological relevance remains largely unknown. ERK is known to target multiple Ser/Thr residues on Gab1 that are predominantly implicated in signal attenuation [41]. Ser552 phosphorylation of Gab1 by ERK was reported to have a positive role in erythropoietin (Epo) signaling [21]. In the context of diverse signaling networks, global phosphoprotein analyses indicate Ser266 phosphorylation in multiple tissues [14,15,31].

The serine/threonine kinase Salt Inducible Kinase SIK 2 (SIK2), together with its paralogs SIK1 and SIK3, belongs to the AMPK superfamily of regulators of energy metabolism [11]. In this regard SIK2 participates in the regulation of hepatic gluconeogenesis and lipogenesis downstream of hormonal control, by modulating the CREB co-activator TORC2 and HAT p300 activities respectively [10,19,54,62]. In adipocytes SIK2 was reported to phosphorylate IRS1 resulting in attenuation of insulin signaling [30], and was also implicated in downregulation of CREB-dependent lipogenic gene expression [50]. Recently, SIK2 was shown to be involved in glucose-induced insulin secretion from pancreatic β-islet cells [58]. Reported non-metabolic functions of SIK2 include regulation of cellular processes as diverse as melanogenesis [29], cardiac hypertrophy [56], endoplasmic reticulum-associated protein degradation (ERAD) [70], autophagosome maturation [71], and macrophage reprogramming [17]. SIK2 activity was also implicated to play a role in cell cycle progression through regulating localization of the centrosome linker protein C-Nap1 [2]. In the CNS, SIK2 is involved in corticotropin-releasing hormone transcription and was associated with survival of hippocampal neurons after ischemia via TORC1 phosphorylation [45]. SIK2 is also abundantly expressed in retinal cells. In Müller glia it downregulates the insulin-dependent PI3K/Akt survival pathway by targeting IRS1 [38]. Phosphorylation of SIK2 on Thr175 by LKB1 is essential for its catalytic activity [11,47,54]. PKA was proposed to negatively regulate SIK2 by targeting Ser343/Ser358 and Thr484/Ser587 [33,64,65]. In the context of gluconeogenesis and lipogenesis, SIK2 is phosphorylated on Ser343, Ser358, Thr484 and Ser587, however there is no consensus on whether these phosphorylation events modulate catalytic activity [19,25,54]. Thr484 phosphorylation by Ca2^+^/calmodulin-dependent protein kinase I/IV was suggested to promote its proteosomal degradation of SIK2 [60]. In HEK293T cells, SIK2 forms a complex with PP2A, which is suggested to contribute to the maintenance of their respective activity [40]. Previously, our lab identified SIK2 as a weak interactor of FGFR2 in a yeast 2 hybrid screening of retinal cDNA library (data not shown), implicating the protein in FGF signaling.

In this study, we present evidence supporting SIK2 involvement in the negative regulation of FGF2 dependent cell proliferation in the context of retinal Müller glia. We propose that ERK-mediated activation of SIK2 targets Gab1, resulting in downregulation of the FGF dependent Ras/ERK cascade in a negative feedback loop. This previously uncharacterized mechanism potentially contributes to prevent excessive signaling through FGFRs.

## Materials and Methods

### Plasmids Constructs and Transfections

The c-Myc tagged human Gab1 cDNA clone (RC209622) was purchased from Origene, USA. Cloning of the human SIK2 coding sequence to the pEGFP-C3 (Clontech, USA) vector was previously described by Küser-Abali et al., 2013 [38]. MIO-M1 cells were transfected with 20 µg/ml of pEGFP or pEGFP-SIK2 vector using FugeneHD (Roche, Germany) according to the manufacturer’s instructions. An IRS1 fragment containing SIK2 target residue was obtained as described before [38]. The GST-IRS1 fusion protein was expressed in E. coli BL21 and purified using glutathione-Sepharose 4B microspin columns (Amersham, USA) following the manufacturer’s instructions.

### Cell Culture

The spontaneously immortalized MIO-M1 Müller glia cell line was kindly provided by Dr. G. A. Limb (University College London, Institute of Ophthalmology). Cells were maintained in DMEM-glutamax (Invitrogen, USA) containing 25 mM glucose supplemented with 10% fetal bovine serum (Biochrom, Germany) and 0.1% penicillin/streptomycin (Invitrogen, USA) under 5% CO2. For FGF2 treatment sub-confluent MIO-M1 cell cultures were serum starved overnight prior to treatment with 1 ng/ml FGF2 (R&D Systems, USA) and 10 μg/ml heparin for indicated times. At the end of incubation periods, cells were washed with ice-cold PBS containing protease and phosphatase inhibitor cocktails (Roche, Germany), scraped and pelleted by centrifugation.

In experiments involving inhibition of SIK2 activity, Compound C (Sigma-Aldrich, USA) was added to the cultures 30 minutes prior to FGF2 treatment. The final concentration of the inhibitor was 0.5 µM where it was reported not to have any effect on AMPKs [60].

### Immunoprecipitation and Co-immunoprecipitation

For immunoprecipitations an SIK2 antiserum kindly donated by Dr. Takemori (National Institute of Biomedical Innovation, Osaka, Japan at 1:125 dilution) and anti-Gab1 antibody (Santa Cruz-H-198, at 1:25 dilution) were used. Briefly, cell lysates were prepared in a buffer containing 50 mM Tris-HCl (pH 8.0), 150 mM NaCl, 1% NP40 supplemented with protease and phosphatase inhibitor cocktails (Roche, Germany) and pre-cleared with protein A-agarose beads (Roche, Germany). Primary antibody incubations were carried out for 2 hours at 4°C, followed by another 2-hour incubation with protein A-agarose beads. Beads were collected with centrifugation, extensively washed with lysis buffer and boiled in SDS-PAGE sample buffer for 5 min. for western blotting. Immunoprecipitation with normal IgG or incubation of lysates with protein A-agarose beads alone constituted the negative controls, and in no case produced signals in the western blots. The absence of the known interacting partners in the immunoprecipitates was verified by western blotting using antibodies against SIK2, Gab1, Grb2 and Shp2.

Co-immunoprecipitations were done essentially the same way except the lysis buffer contained 50 mM Tris-Cl (pH 8.0), 300 mM NaCl, 5 mM EDTA, 5 mM EGTA, 2 mM DTT and 0.5% TritonX-100 with protease and phosphatase inhibitor cocktail. After the incubation, the beads were washed twice with ice-cold IP buffer (20 mM HEPES (pH 7.4), 150 mM NaCl, 0.1% Triton-X and 10% glycerol) with protease and phosphatase inhibitor cocktail and once with PBS.

### *In vitro* Kinase Assay

To evaluate kinase activity *in vitro*, SIK2 immunoprecipitated from lysates prepared from 10^7^ cells and resuspended in the kinase buffer containing 50 mM Tris-Cl 9 (pH 7.4), 1 mM DTT, 10 mM MgCl2, 10 mM MnCl2. Half of the samples were analyzed by western blotting to assess the inputs. The remaining samples were incubated with 1 µCi [γ32P]-ATP (3000 Ci/mmol; Isotop, Hungary) and 500 ng of purified recombinant GST-IRS1 at 30^0^C for 1 hour, the reactions were terminated by the addition of the SDS sample buffer and heated to 95^0^C for 5 min. The beads were removed by centrifugation, the supernatants were fractionated on 8% SDS-PAGE. The gels were fixed, dried and exposed to X-ray film (Amersham, UK) for varying times at -80^0^C.

To investigate whether ERK2 directly phosphorylates SIK2, constitutively active ERK2 (R&D Systems) and immunoprecipitated GFP-SIK2 (KI) proteins were included in the kinase reaction. To assess the inhibitory activity of Compound C on SIK2 The Universal Kinase Activity Kit (R&D, USA) was used as instructed by the manufacturer. Briefly, immunoprecipitated SIK2 was incubated in a buffer containing 25 mM HEPES pH 7.0, 0.15 M NaCl, 10 mM MgCl2, 10 mM CaCl2, 0.2 mM ATP and 2 ng/ ml CD39L phosphatase for 30 minutes at room temperature. Subsequently, the reaction was stopped using the malachite green containing reagents supplied by the manufacturer. The amount of inorganic phosphate released by the coupling phosphatase was measured at 620 nm, kinase activity was calculated by referencing a phosphate standard curve . The calculated activity were normalized to the SIK2 levels in the same samples as determined by western blotting.

### Western Blotting

Samples fractionated on polyacrylamide gels were electroblotted to polyvinyl difluoride (PVDF) membranes (Roche, Germany) as described before [16,38]. Following blocking with 5% skimmed milk powder in 150 mM NaCl, 20 mM Tris-HCl (pH 8.0), 0.1% Tween 20 for 1 hour at room temperature, the membranes were incubated with one of the primary antibodies. Subsequently, membranes were washed, incubated with appropriate HRP-conjugated secondary antibodies, and bands were visualized using Lumi-light Western blotting substrate (Santa Cruz Biotechnology, Santa Cruz, CA) for 1 minute, and images were captured using Stella gel imaging system (Raytest, Germany).

Polyclonal SIK2 antibody (rabbit, Novus Biologicals, USA) was used at 1:2500 dilution, anti-GFP (Abcam, UK) at 1:1000, pERK (Santa Cruz Biotechnology, Santa Cruz, CA) and ERK2 (Santa Cruz Biotechnology, Santa Cruz, CA), pTyr (Santa Cruz Biotechnology, Santa Cruz, CA), pSer (Zymed, USA), pThr (Zymed, USA), Gab1 (Santa Cruz Biotechnology, CA), Grb2 (Upstate, USA), Shp2 (Santa Cruz Biotechnlogy, CA), β-Actin (Santa Cruz Biotechnology, CA), FGFR2 (Santa Cruz Biotechnology, CA) antibodies were used at 1:1000 dilution.

### SIK2 Gene Silencing

Human SIK2 sh-RNA lentiviral particles, a pool of three target-specific constructs, and scrambled sh-RNA were purchased from Santa Cruz (USA). MIO-M1 cells were subjected to lentivirus infection as instructed by the manufacturer and propagated under puromycine (1mg/ml, Sigma, USA) selection. After 2 weeks colonies were isolated and analyzed for SIK2 downregulation. Positive colonies were stored at -80^0^C until further use.

### Cell Proliferation Assay

Stably transduced sh-SIK2 or scrambled sh-RNA containing MIO-M1 cells were treated with 1 ng/ml or 100 ng/ml FGF2 containing 10 µg/ml heparin or vehicle (10 µg/ml heparin in PBS) for 24 hours. To assess proliferation, 10 mM BrdU (Roche, Germany) was added to the culture medium, incubated for 5 hrs and analyzed by anti-BrdU immunocytochemistry. To visualize the nuclei, cells were incubated with DAPI for 5 minutes. The samples from three independent experiments were evaluated by fluorescent microscopy. DAPI labeled nuclei were scored by counting a minimum of 200 cells per sample.

### Site-directed Mutagenesis

To create a kinase inactive mutant of human SIK2 (SIK2-K49M), lysine 49 residue was replaced with methionine by overlap extension PCR using the primer pair: 5’-CCAAGACGGAGGTGGCAATAATGATAATCGATAAGTCTCAGC-3’ and 5’-GCTGAGACTTATCGATTATCATTATTGCCACCTCCGTCTTGG-3’. Human Gab1 serine 266 was replaced with alanine (Gab1-S266A) using the primer pair: 5’-CTGCCCAGGAGTTATGCCCATGATGTTTTAC-3’ and 5’-

GTAAAACATCATGGGCATAACTCCTGGGCAG-3’. For the site-directed mutagenesis reactions the QuickSiteXL Mutagenesis kit (Stratagene, USA) was used and the mutant were verified by sequencing.

### Statistical analysis

Experimental groups were compared statistically using the Mann-Whitney test (one-tailed). Means with p<0.05 were considered statistically significant.

## Results

### SIK2 Phosphorylation/Activity are Modulated by FGF2 Stimulation and SIK2 Interacts with FGFR2 *in vivo*

A number of studies indicated that serine and threonine phosphorylation of SIK family members are important in regulation of their kinase activity in response to various extracellular stimuli [7,19,24,27,47,57,68] . Therefore, possible FGF2-dependent modulations in serine and threonine phosphorylation of SIK2 were assayed using SIK2 immunoprecipitated from growth factor treated (0-60 min) cells and subjected to Western blotting with antibodies specific to pSer and pThr. When pSer and pThr band intensities were normalized to that of the input SIK2 (Figure 1A), a modest decline in serine phosphorylated SIK2 levels were detectable at 5 minutes post induction, the minimum levels, representing a 50% decrease, were reached at 10 minutes and basal levels were restored by 60 minutes of induction. Threonine phosphorylated SIK2 levels, on the other hand, increased within 5 minutes of FGF2 treatment, peaked at 65% increase over resting cells at 10 minutes of induction, and remained about the same thereafter. At no time point examined did we observe signal with the two different anti-phosphotyrosine antibodies that were tested (data not shown).

**Figure 1.**
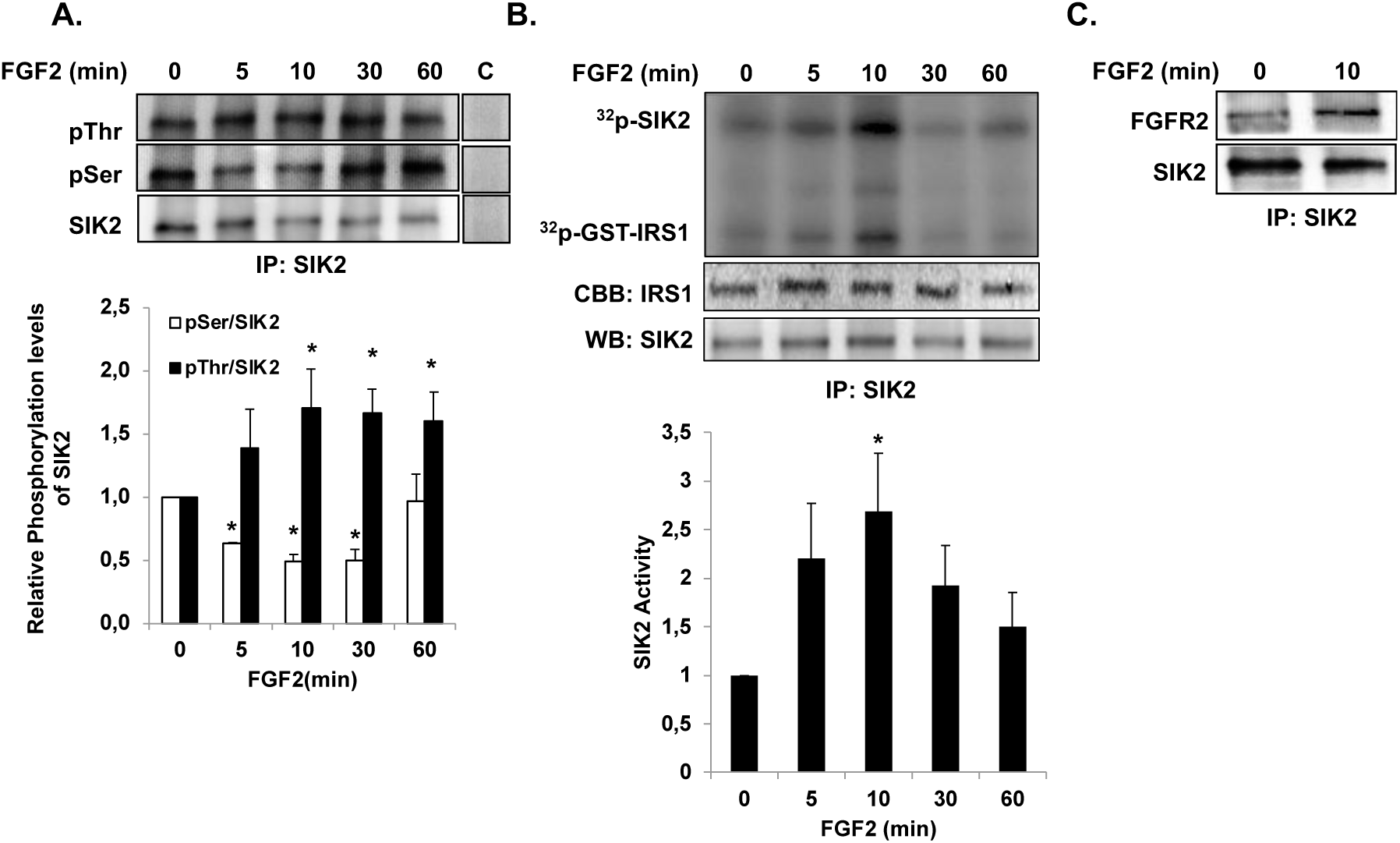
FGF2-dependent changes in phosphorylation status/ activity of SIK2 and SIK2/FGFR2 interaction. Lysates from MIO-M1 cells were treated with FGF2 were subjected to immunoprecipitation with anti-SIK2 antibody (A) Immunoprecipitated material was immunoblotted with anti-pThr, anti-pSer or anti-SIK2 antibodies, in lane C antibodies were pre-incubated with respective blocking peptides. Graphic representation of SIK2 phosphorylation levels normalized to that of corresponding SIK2 is shown in the lower panel. (B) GST-IRS1 phosphorylation by immunoprecipitated SIK2 was assayed in *vitro* in the presence of ^32^P-γ-ATP, followed by PAGE and autoradiography. The input levels of SIK2 were revealed by Western blot analysis using anti-SIK2 antibody and by Coomassie blue staining of GST-IRS1. Graphic representation of pGST-IRS1 band intensities normalized to that of SIK2 levels in the same samples is shown in the lower panel. C. SIK2 immunoprecipitated complexes obtained from cells treated with FGF2 for 10 min and the unstimulated control cells were subjected to immunoblotting with anti-FGFR2. Three independent experiments were carried out in each case. * *P* < 0.05, compared to corresponding untreated samples.

To investigate whether SIK2 activity was modulated in an FGF2-dependent manner in this time frame, in vitro kinase assays were performed using immunoprecipitated SIK2 in the presence of GST-IRS1 and ^32^P-γATP. When densitometric readings of GST-IRS1 bands on autoradiograms were normalized to inputs, SIK2 activity increased nearly 3 fold within 10 minutes of FGF2 induction and gradually returned to basal levels within 60 minutes (Figure 1B).

Next we examined if endogenous SIK2 interacts with FGFR2 by coimmunoprecipitation using serum starved MIO-M1 cells treated with FGF2 for 10 minutes where SIK2 activity is at maximum, and for the controls cells were left untreated. Evaluation of SIK2 immunoprecipitates for the presence of FGFR2 revealed a low level of signal in the untreated samples. The amount of FGFR2 co-immunoprecipitated with SIK2 was doubled upon FGF2 induction (Figure 1C). The SIK2/FGFR2 interaction revealed here agrees with our yeast-two-hybrid data.

Taken together these findings support the involvement of SIK2 in the FGF2 signaling. It is conceivable that concurrent threonine hyperphosphorylation and serine hypophosphorylation of SIK2 in response to FGF2 lead to a rapid increase in the kinase activity of the protein. As phosphoserine levels return to baseline levels after 30 minutes of FGF2 induction, SIK2 activity also declines. Because no tyrosine phosphorylation of SIK2 could be detected, SIK2 is not likely to be a direct target of FGFR.

### SIK2 negatively regulates FGF2-dependent ERK activation and Müller cell proliferation

MIO-M1 cells proliferate in response to FGF2 via ERK activation [28]. Therefore, we investigated the effect of changes in SIK2 expression levels on FGF2-dependent pERK levels by immunoblotting and on MIO-M1 proliferation by BrdU incorporation assays.

In wild-type MIO-M1 cells ERK activation exhibited a transient profile, reaching peak levels (3.6 fold of basal level) within 10 minutes of FGF2 treatment and thereafter a gradual decrease was observed (Figure 2A), similar to cultured primary Müller glia (unpublished data).

**Figure 2.**
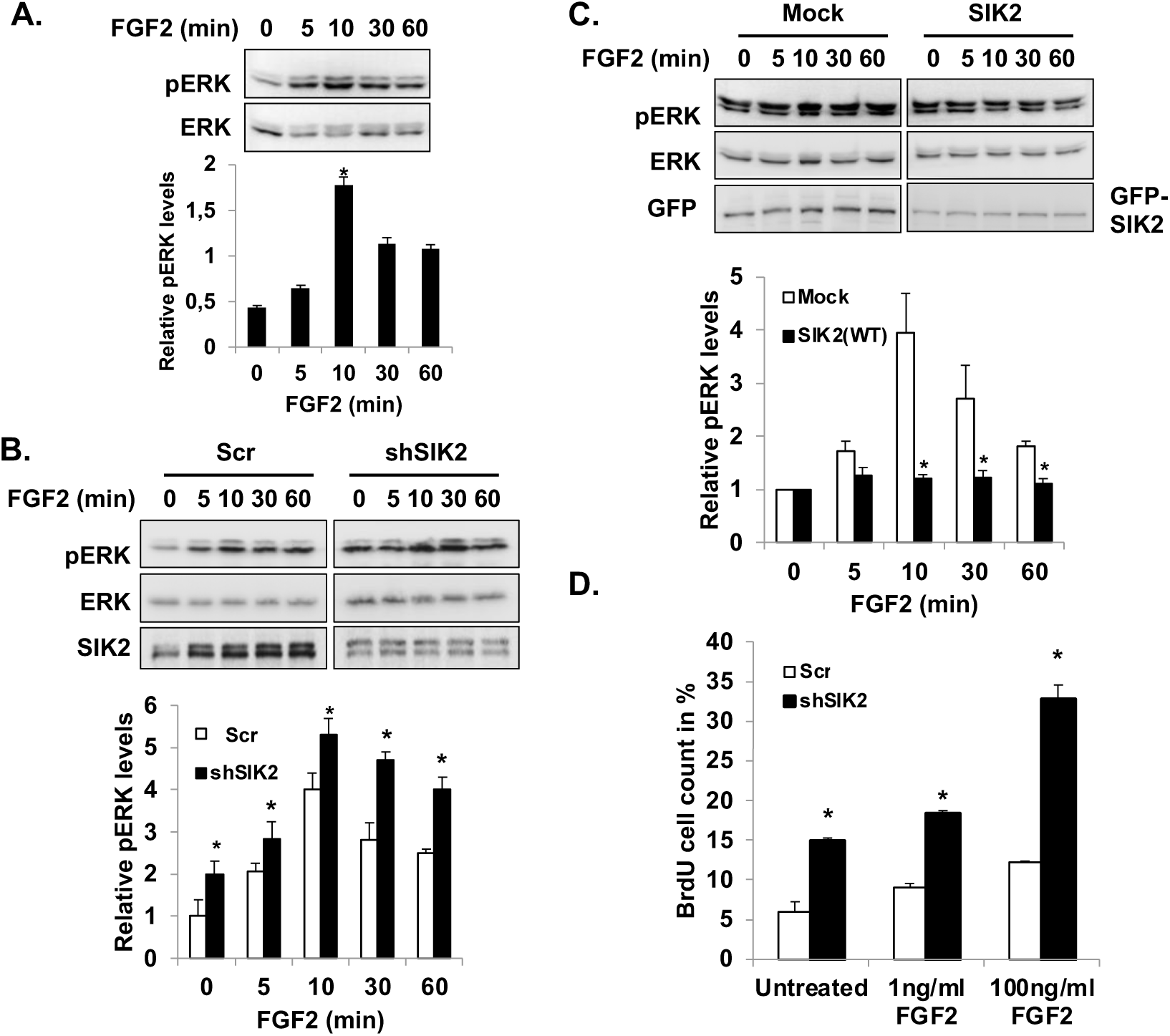
Effect of SIK2 on FGF2-dependent ERK activation and proliferation. MIO-M1 cells were treated with FGF2 for the indicated times. Active ERK levels were assessed by Western blot analysis with anti-pERK, and total ERK by probing the membranes with anti-ERK antibodies; the pERK band intensities were normalized to that of ERK in the same samples and represented as mean ±SE of three independent experiments in each case. (A) Western blots were done using wild-type MIO-M1 cell lysates. (B) MIO-M1 cells transfected with either pEGFP-SIK2 or only the vector received FGF2 treatment 2 days post-transfection. (C) Cells were stably transduced with lentiviral particles carrying either sh-SIK2 or scrambled (Scr) sh-RNA. (D) Proliferation was evaluated by BrdU assay using stably transduced cells that were serum-starved overnight, subsequently treated with 1 or 100 ng/ml FGF2 for 24 hours, control cells did not receive growth factor induction. Data represent the mean ±SE of percent BrdU positive cells from three independent experiments. In all cases \**P* < 0.05 compared to the corresponding untreated samples. n.s.: non-significant.

For enhanced SIK2 expression studies, cells were transfected with pEGFP-SIK2 plasmid at 40% transfection efficiency. In mock transfected cells, the ERK phosphorylation profile showed a transient nature as the wild-type MIO-M1 cells, peaking to 4-fold at 10 min of FGF2 stimulation (Figure 2A and Figure 2B). In contrast, SIK2 overexpression did not result in significant upregulation of ERK activation over the basal level at any time points examined (Figure 2B).

When SIK2 protein levels were downregulated by about 60% in sh-SIK2 transfected cells, pERK levels were 1.5-2 fold higher than the cells transfected with scrambled sh-RNA at all time points examined, including the resting cells. The fraction of active ERK increased at comparable rates for both groups during the first 10 min of growth factor treatment (Figure 2C), but a prolonged ERK activation was observed in sh-SIK2 transfected cells. In control samples a 30% decrease in active ERK levels was evident after 30 min of FGF2 treatment similar to untransfected cells. However, at this time point there was no statistically significant reduction of pERK levels could be detected in the SIK2 depleted cells.

In the presence of Compound C, a relatively selective chemical inhibitor of SIKs, significant reduction in SIK2 autophosphorylation activity was evident (Figure 2D). We observed no downregulation of FGF-dependent ERK activation in the time frame of the experiment.

Downregulation of SIK2 by shRNA also resulted in an increase in MIO-M1 cell proliferation. Under serum-starvation, the fraction of BrdU positive cells was 3 fold higher in SIK2 downregulated samples compared to the control cells transfected with scramble shRNA (Figure 2E). Upon FGF2 treatment both groups responded with an increase in cell proliferation in a concentration dependent manner, and again with SIK2 silencing BrdU labeled cell numbers were 2-3 fold higher than that of the control group.

Overall these data suggest that SIK2 is a negative regulator of FGF2-dependent ERK activation and proliferation of MIO-M1 cells.

### SIK2 phosphorylates Gab1 docking protein on serine 266

Scanning of the known RTK pathway elements for the presence of the consensus SIK2 phosphorylation motif, L[(B)X or X(B)][S/T]XSXXXL, [30,62] identified LPRSYS**^266^**HDVL sequence in Gab1 as a potential target site. Gab1 is one of the key regulatory proteins in RTK signaling [26,36,37,52], Gab1 serine phosphorylations have been associated with reduced signal relay [23,72]. A putative serine phosphorylation residue at position 266 and its surrounding sequence are strictly conserved among the different mammalian species (http://www.ncbi.nlm.nih.gov/homologene?cmd=Retrieve&dopt=MultipleAlignment &list_uids=1542). Thus, we hypothesized that SIK2 modulates FGF2-dependent ERK activation via phoshorylation of Ser266 on Gab1.

The possibility of Gab1 being an SIK2 substrate was investigated by in vitro kinase experiments using immunopurified GFP–SIK2 and Myc–Gab1 recombinant proteins expressed in MIO-M1 cells in the presence of ^32^P-γ-ATP. The autoradiograms of the reaction products fractionated on SDS-PAGE revealed that SIK2 is indeed capable of catalyzing Gab1 phosphorylation (Figure 3A, lane 1). No signal was detected when kinase-defective mutant SIK2 (K49M) (Figure 3A, lane 3) or FRS2 adaptor lacking the canonical SIK2 target motif included as the substrate (data not shown). Further, the use of a mutant Gab1 (S266A) in the kinase assay completely eliminated the phosphorylation signal corroborating the specificity of SIK2 phosphorylation at this residue (Figure 3A lane 2).

**Figure 3.**
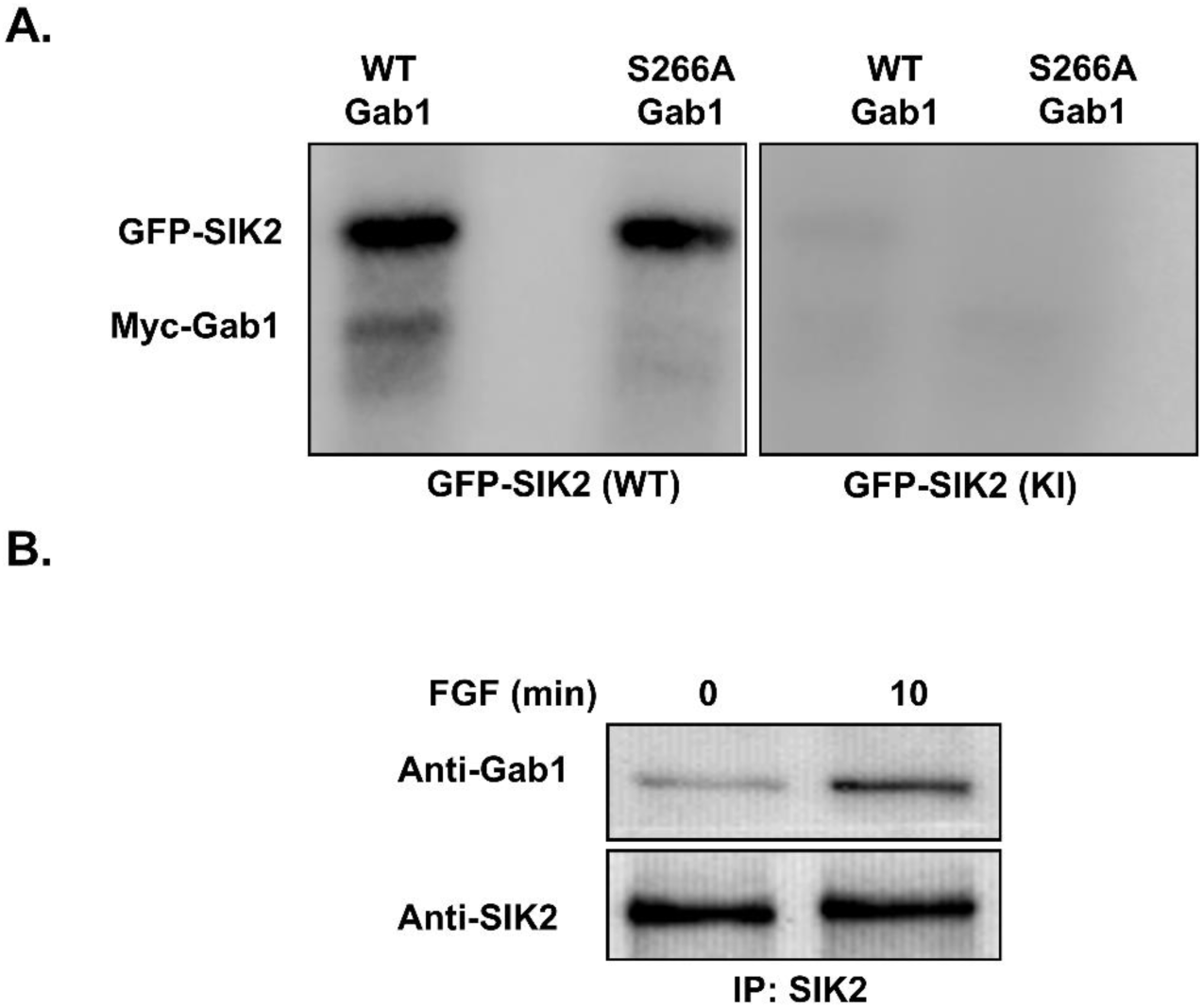
Gab1 Ser266 phosphorylation by SIK2 and Gab1-SIK2 interaction. (A) GFP-tagged wild-type and kinase dead SIK2 (K49M), Myc tagged wild-type and S266A mutant Gab1 were expressed as fusion proteins in MIO-M1 cells, and were purified using anti-tag antibodies. In *in vitro* kinase assays Gab1 fusion proteins were used as substrates either with wild-type *(left panel)* or kinase dead SIK2 *(right panel)* in the presence of [γ^32^P]-ATP. The reaction mixtures were fractionated by SDS PAGE and subjected to autoradiography, the presence of SIK2 activity was assessed by autophosphorylation signal. (B) SIK2 immunoprecipitations using anti-SIK2 were carried out using the lysates from MIO-M1 cells to reveal interaction of endogenous proteins. Cells were either untreated or treated with FGF2 for 10 min, Western Blot analysis were done using either anti-SIK2 or anti-Gab1 antibody.

To explore the possible SIK2 and Gab1 interaction *in vivo*, MIO-M1 cell lysates were subjected to co-immunoprecipitation with anti-SIK2 antibody followed by Gab1 western blotting. In serum-starved cells SIK2 and Gab1 interact at low levels, this interaction was significantly enhanced following 10 min of FGF2 treatment where the SIK2 activity is at maximum (Figure 3B). Reciprocally, SIK2 was pulled down when anti-Gab1 antibody was used for immunoprecipitation (data not shown).

These data are consistent with the hypothesis that SIK2 may regulate FGF2 signal transduction via phophorylation of Gab1 at Ser266 in MIO-M1 cells.

### Silencing of SIK2 gene expression and S266A mutation of Gab1 increases interactions of Gab1 with docking partners

In RTK signaling Gab1 becomes tyrosine phosphorylated and recruits various signal relay molecules, the most prominent ones being Grb2, Shp2 and p85 [63]. Gab1/Grb2 and Gab1/Shp2 associations were shown to promote Ras/ERK pathway activation, whereas Gab1-p85 binding is directly involved in PI3K/Akt pathway activation [26,52].

In the initial experiments Gab1 was immunoprecipitated from cells treated with FGF2 and subjected to immunoblotting with anti-pSer, anti-pTyr or anti-Grb2 antibodies. The results showed global increase in phosphorylation on tyrosine and serine residues within 10 minutes of FGF2 treatment, which then returned to basal levels by 60 minutes (Figure 4A). Coimmunoprecipitation of Grb2 and Gab1 also reached maximum at 10 minutes after growth factor induction and declined to basal level by 30 minutes (Figure 4A). These observations verify Gab1 involvement in the relay of FGF signal in Müller cells.

**Figure 4.**
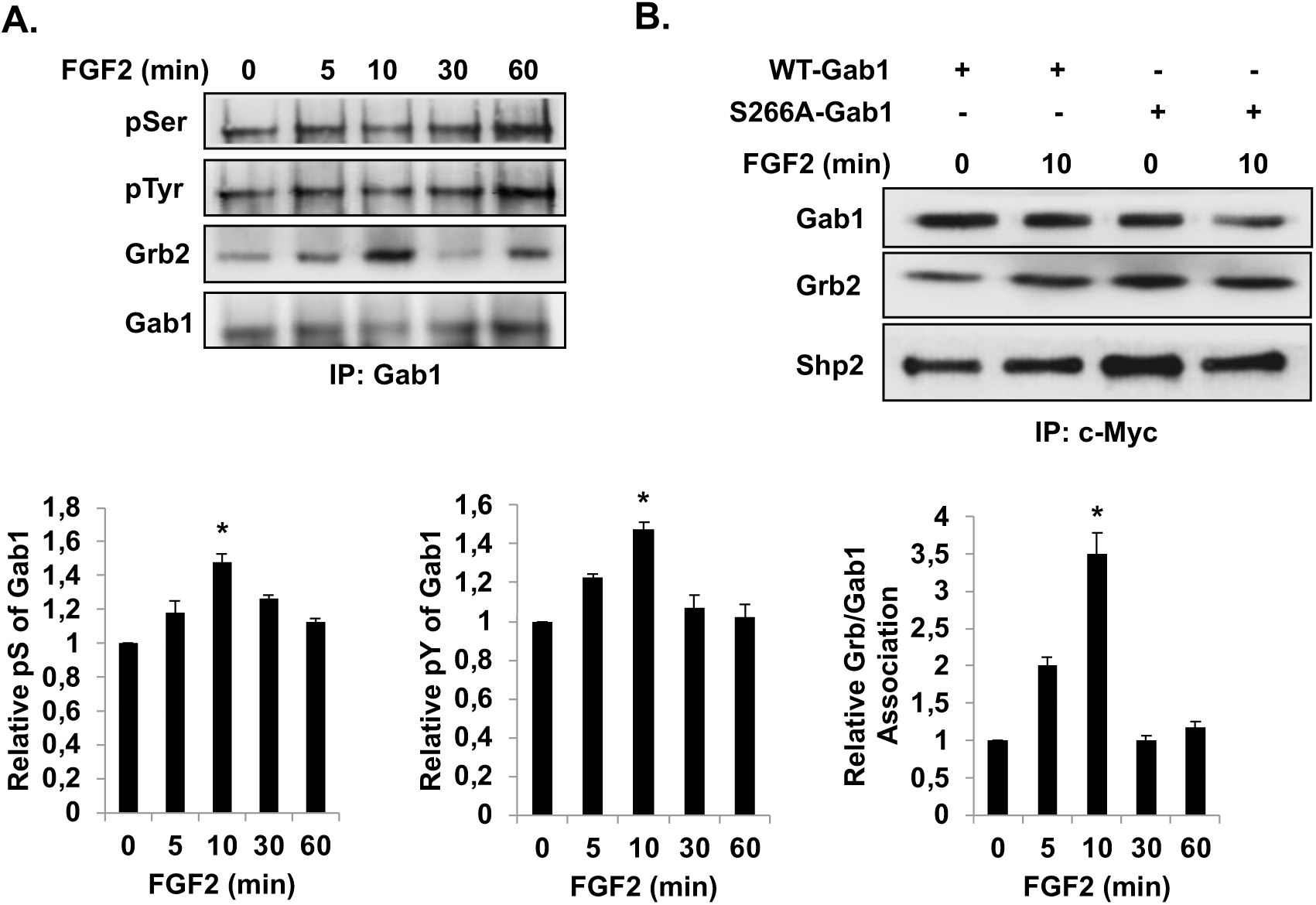
FGF2-dependent Gab1 phosphorylation and its association with the partners. (A) Gab1 was immunoprecipitated by using anti-Gab1 antibody from lysates of MIO-M1 cells treated with FGF2 for indicated times subsequent to overnight serum starvation. Immunoprecipitates were analyzed by Western blotting using anti-pSer and anti-pTyr, anti-Grb2 and anti-Gab1 antibodies as indicated. The band intensities normalized to that of Gab1 in the same samples, mean±SE of three independent experiments are represented in the graphs. * *P* < 0.05 compared to the untreated samples. (B) MIO-M1 cells overexpressing Myc tagged WT-Gab1 or S266A-Gab1 were untreated/treated with FGF2 for 10 minutes. Immunoprecipitations were carried out with anti-Myc antibody, subsequently the samples were subjected to western blot analysis with anti-Myc, anti-Shp2 and anti-Grb2 antibodies.

We next transfected MIO-M1 cells with myc-tagged wild-type Gab1 or S266A mutant protein, and treated the cells with FGF2 for 10 minutes. Immunoprecipitation with anti-Myc antibody and subsequent immunoblotting with anti-Grb2 and anti-Shp2 antibodies showed that mutant Gab1 exhibited significantly enhanced binding to Grb2 and Shp2, compared to wild-type Gab1 (Figure 4B). These experiments suggest that Ser266 phosphorylation may weaken Gab1 interaction with its partners.

The downregulation of SIK2 expression by shRNA results in 40% decrease in the overall levels of serine phosphorylated Gab1, but increases the levels of tyrosine phosphorylated Gab1 3-fold when compared to the control cells carrying scrambled shRNA (Figure 5A). Furthermore, SIK2 depletion enhanced Gab1 interaction with Grb2 and Shp2, 2 and 1.5-fold, respectively. It should be noted, however, that SIK2 silencing increased Gab1 protein level by 90% (Figure 5B).

**Figure 5.**
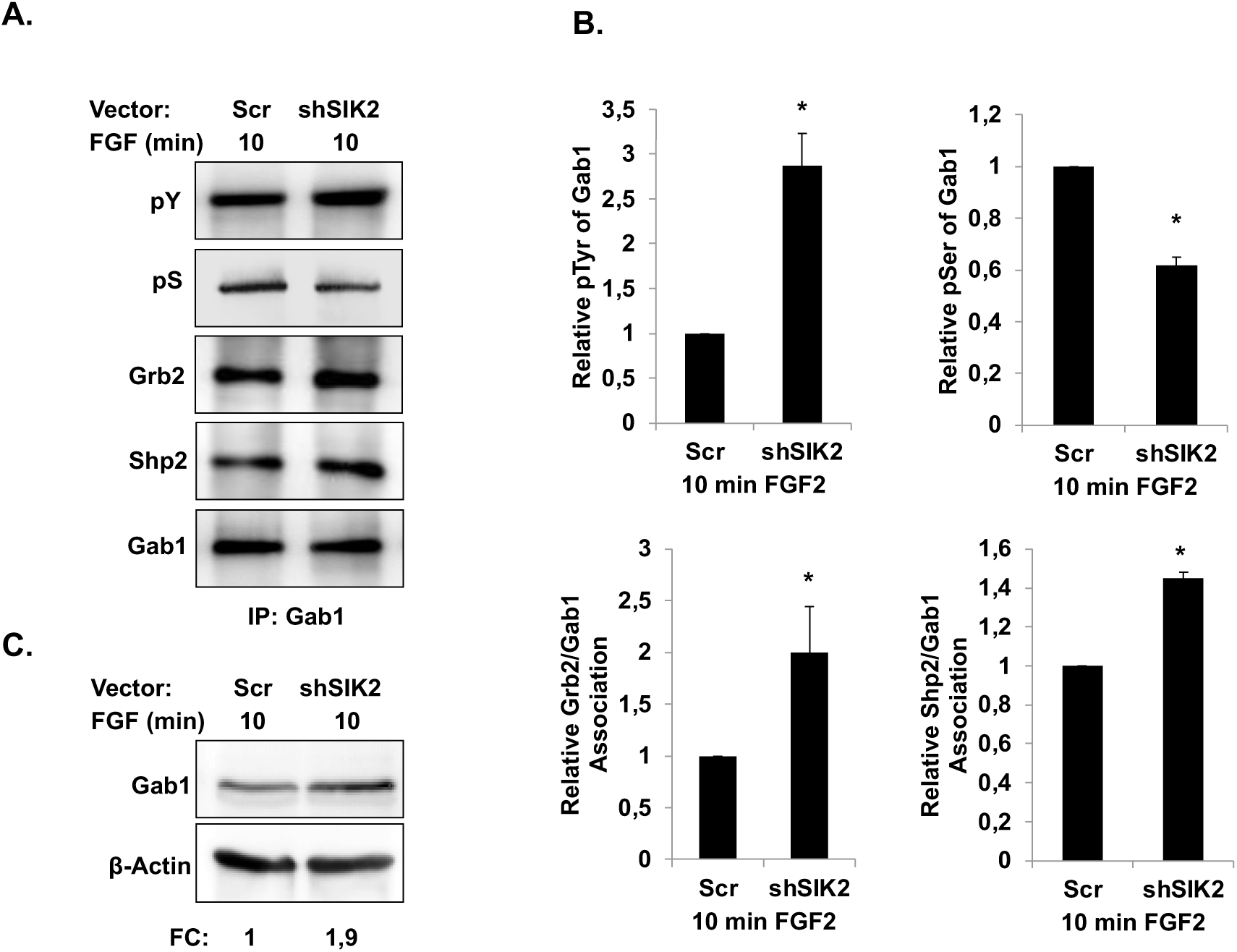
Effect of SIK2 gene silencing on FGF2-dependent Gab1 phosphorylation and binding partner interactions. MIO-M1 cells stably transduced with scrambled RNA or shSIK2 were serum starved overnight, then treated or not with FGF2 for 10 minutes. (A) Gab1 was immunoprecipitated using anti-Gab1 antibody. Immune complexes were subjected to Western blot analysis using anti-Gab1, anti-pSer and anti-pTyr, anti-Grb2 or anti-Shp2 antibodies. Graphic representation of indicated band intensities normalized to that of Gab1 in the same samples. Data represent the mean±SE of three independent experiments. * p < 0.05 compared to the control samples. (B) Equal amounts of proteins from cells treated as in panel A were analyzed for endogeneous Gab1 levels by probing the immunoblots with anti-Gab1 antibody, anti-β-actin probing indicates input levels. FC: Fold change

These findings suggest that SIK2 exerts its negative effect on FGF2 signal transduction by phosphorylating the target residue Ser266 on Gab1, which may hamper Gab1 tyrosine phosphorylation and weakens Gab1’s ability to bind to the docking partners. The reduced Gab1 protein levels may also contribute to the observed decrease in Gab1 interaction with Grb2 and Shp2.

### ERK is an upstream kinase of SIK2 and modulates its activity upon FGF2 stimulation

Scanning of the SIK2 sequence revealed that it has potential ERK phosphorylation, (S/T)P and FxFP (aa: 399-402), and DEJL (aa: 605-616) docking motifs. Since in the regulation of FGF2 pathway feedback loops of ERK are important [39], we explored whether ERK phosphorylates threonine residues and thereby activates SIK2. To eliminate autophosphorylation, a GFP-tagged kinase-inactive (SIK2 K49M) mutant form of SIK2 was expressed in MIO-M1 cells and immunopurified with anti-GFP antibody, and subsequently was used in *in vitro* kinase assays in the presence of constitutively-active ERK. When only SIK2K49M was included in the\ reaction no phosphorylation signal was detectable (Figure 6A), but SIK2 was phosphorylated strongly when ERK was present.

**Figure 6.**
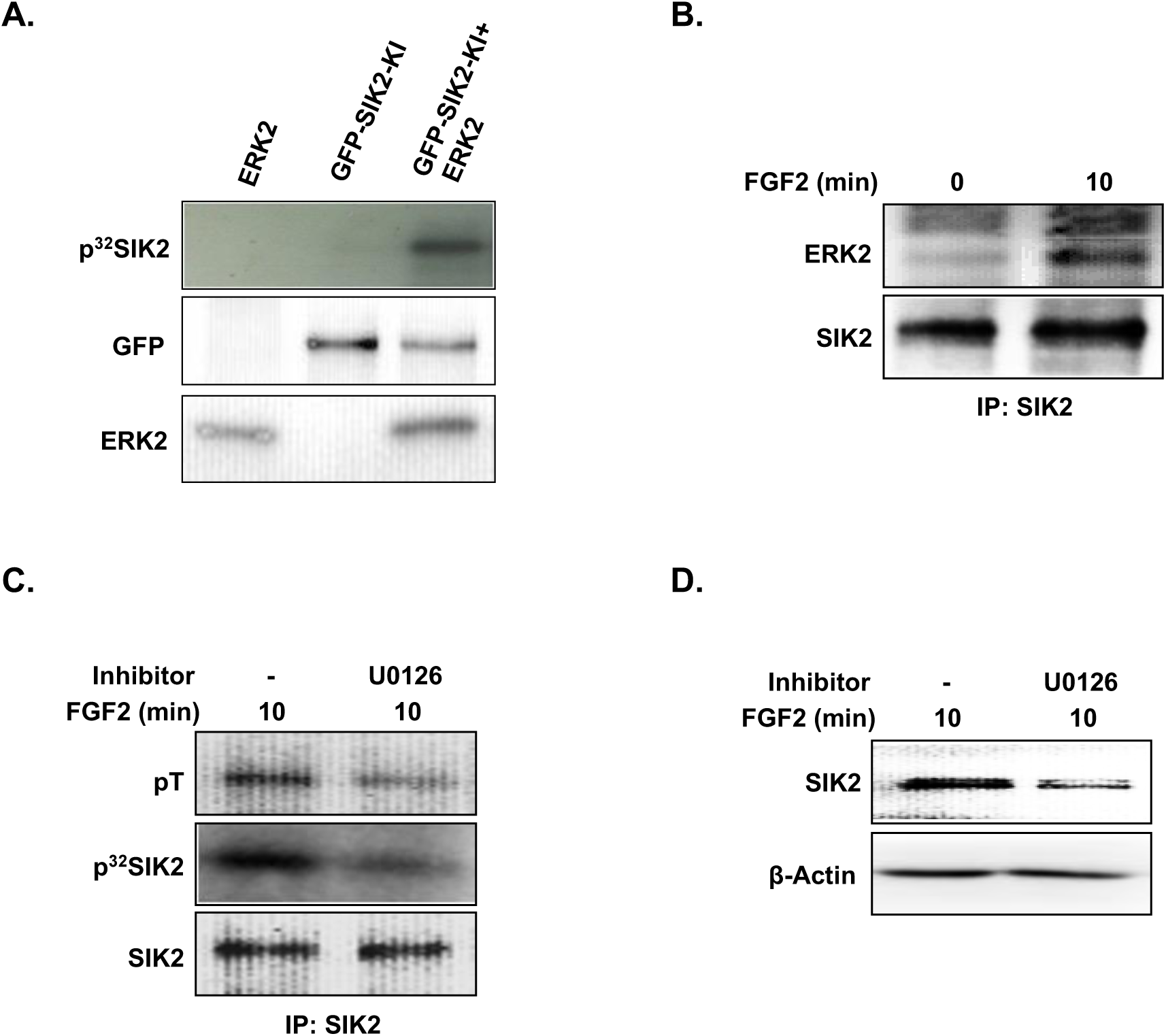
ERK as the upstream activatory kinase of SIK2. (A) *In vitro* kinase assays were carried out using constitutively active recombinant ERK2 and kinase inactive GFP-SIK2; in controls either kinase dead SIK2 or the ERK was present (first two lanes). Phosphorylation signal was detected by autoradiography, Western blots probed with GFP or ERK2 indicate the protein input levels. (B) SIK2 immunoprecipitations were carried out using the lysates obtained from MIO-M1 cells transfected with wild-type SIK2 that were untreated or treated with FGF2 for 10 min, Western Blots were probed with either anti-ERK1/2 or anti-SIK2 antibodies. (C) Serum-starved MIO-M1 cells treated 30 min with MEK inhibitor U0126 prior to 10 minute FGF2 induction; samples treated only with FGF2 for 10 minutes constituted the positive control. Subsequently SIK2 was immunoprecipitated followed by Western Blot analysis with anti-pThr antibody (top panel); and used to assess autophosphorylation *in vitro,* followed by PAGE and autoradiography *(middle panel)*. Input SIK2 was shown by Western blot analysis using anti-SIK2 antibody *(lower panel)*. (D) Lysates from cells treated as in panel C were analyzed for endogenous SIK2 levels by probing the immunoblots with anti-SIK2 antibody, anti-β-actin antibody probing indicates input levels.

Coimmunoprecipitation of endogenous SIK2 revealed interation with ERK and that this interaction is enhanced upon FGF2 treatment (Fig 6B). To investigate whether ERK is involved in threonine phosphorylation and activation of SIK2, cells were treated with MEK inhibitor U0126 prior to 10 minutes of FGF2 exposure. Immunoprecipitated SIK2 was then subjected to immunoblotting and evaluated by in vitro kinase assays. Probing of the blots with anti-phosphothreonine and SIK2 antibodies showed that U0126 treatment prior to FGF2 stimulation results in a 50% reduction in both threonine phosphorylation and overall SIK2 levels (Fig 6C and D). Furthermore, the activity of SIK2 was half the levels of the cells that were not exposed to MEK inhibition.

These findings suggest that ERK may be an upstream kinase in FGF2-dependent SIK2 threonine phosphorylation and upregulation of its activity and/or stabilization. Altogether our results support the model that SIK2 is an element of a negative feedback loop involved in the regulation of the FGF2 signaling pathway.

## Discussion

The possibility of SIK2 being a part of the FGF signaling pathway was raised by its identification as a weak interactor of FGFR2 in a yeast two-hybrid screen (unpublished data). The results presented here pertaining to the rapid and transient changes in SIK2 phosphorylation status and activity following FGF stimulation, and hampered FGF-dependent ERK activation in the presence of increased SIK2 expression levels support this possibility. In the same vein, we observed that downregulation of SIK2 results in increased ERK activation with delayed attenuation, and increased cell proliferation. The ERK activation profile obtained in the presence of compound C at concentrations reported to inhibit SIKs but not the other AMPK family members lends support that the observed delay in ERK activation is dependent on kinase activity of SIK2.

Scanning of the known FGF pathway elements for the SIK2 phosphorylation motif indicated the docking protein Gab1 as a strong potential target for this kinase. Indeed in vitro kinase assays and in vitro mutagenesis studies verified phosphorylation of Gab1 by SIK2 and identified Ser266 as the target residue.

Gab1, as a component of the complex assembled on the activated receptor, takes part in the initiation of mitogenic signaling via the ERK cascade and it is also crucial in the activation of the PI3K/Akt axis promoting survival in response to various growth factors and cytokines [22,44,51,69]. It is generally accepted that Gab1 tyrosine phosphorylation by activated RTKs or PTKs positively modulates its partner interactions [4,5,67], with Grb2 and Shp2 being the most predominant partners in the initiation of the ERK cascade. The interaction with Grb2 is important for the recruitment of Gab1 to the vicinity of RTKs and its subsequent tyrosine phosphorylation and allows Gab1-Shp2 association [18,26,74]. It has been reported that inhibition of Shp2 association with Gab1 and FRS2 decreases FGF2 induced ERK activation in chondrocytes [37]. S/T phosphorylation, on the other hand, is often correlated with reduced tyrosine phosphorylation in docking proteins and implicated in negative regulation of the downstream pathways [9,23]. ERK remains to be the best described upstream kinase of Gab1 with diverse influences on its association with p85 [23,41,72,73]. In insulin signaling the residues Thr312, Ser381, Ser454, Thr476, Ser597 were identified as ERK targets and proposed to interfere with Gab1 tyrosine phosphorylation [41].

We observed a rapid increase in tyrosine phosphorylated Gab1 upon FGF stimulation and enhanced association with its partners Grb2 and Shp2 in Müller cell context. Our studies indicate with SIK2 depletion or with exogenous expression of Gab1 mutated at the putative SIK2 target residue tyrosine phosphorylated Gab1 levels and Gab1 partner interaction are upregulated. It is conceivable that S266 phosphorylation by SIK2 may trigger conformational change that would block activating tyrosine phosphorylation of Gab1 and/or renders the structure of docking platform unfavorable for partner interactions, as proposed earlier for the regulation of different docking proteins [9,23]. This scenario by itself doesn’t account for increased Gab1 levels observed with SIK2 downregulation. Presence of two Crk binding sites found in very close proximity to Gab1 S266 suggests that SIK2 mediated Gab1 phosphorylation at this site may facilitate the binding of Crk and recruitment of CBL (E3Ligase) to promote Gab1 degradation. Therefore, it is possible that SIK2 mediated S266 phosphorylation, in mutually non-exclusive manner, modulates Gab1 availability and interferes with the assembly and/or stability of the signaling complex to promote/amplify ERK activation.

Phosphorylation of Gab1 on Ser266 has been documented in global phosphoprotein analyses of various tissues [31], and in multiple signaling networks [14,15,61], but neither the upstream kinases nor its biological relevance have been described so far. To the best of our knowledge SIK2 is the first enzyme reported to target Gab1 phosphorylation at Ser266. Our results suggest that phosphorylation of S266 hampers Gab1 association with Grb2 and Shp2 and potentially and can be directly correlated to the abrogation of FGF-mediated proliferation. It should be noted this study does not differentiate whether Gab1-Shp2 association is direct or indirect through FRS2-Grb2 complex.

It has been suggested that Gab1 involvement in ERK activation is dependent on PI3K in a signal strength or intensity-dependent manner in different contexts [1,34,59]. Our unpublished data indicate that maximal FGF-dependent p85-Gab1 association and Akt activation were delayed in relation to ERK activation (60 min vs. 10 min post induction). In our experimental model however, ERK activation appears to be independent of the PI3K/Akt branch, although it is possible that basal PI3K activity is required for the initial membrane recruitment of Gab1.

A number of S/T phosphorylation sites on SIK2 have been reported, but their impact on modulation of its activity remains ill defined. It is well accepted that Thr175 phosphorylation by LKB1 in the kinase domain is required for SIK2 activation [11,47,54]. On the other hand, PKA-dependent phosphorylation of SIK2 at Ser343/Ser358 and Thr484/Ser587 generates two pairs of 14-3-3 binding sites and results in negative modulation of its activity [33,64,65]. In the context of TORC2/CREB dependent regulation of gluconeogenesis and lipogenesis, SIK2 phosphorylation on Ser343, Ser358, Thr484 and Ser587 have been detected, however there is no consensus on whether they affect SIK2 catalytic activity [19,25,54]. Recently Thr484 phosphorylation by Ca2+/calmodulin-dependent protein kinase I/IV was suggested to target SIK2 for the proteosomal degradation [60]. We have shown that maximal FGF-dependent SIK2 activity coincides with the concomitant decrease in serine and increase in threonine phosphorylation. Scanning of the SIK2 sequence reveals several potential ERK target and docking motifs. Consistent with this, we have shown that ERK is capable of phosphorylating SIK2 in vitro. Furthermore, we observed coimmunoprecipitation of the endogenous proteins in an FGF-dependent manner, and there is a significant reduction both in SIK2 threonine phosphorylation and activity when ERK activation is inhibited. Taken together these findings suggest that ERK acts as a potential upstream activating kinase of SIK2 in a negative feedback loop controlling the proliferative process. We are currently carrying out studies to identify the specific residues on SIK2 that are phosphorylated by ERK in an FGF2-dependent manner and their potential impact on the catalytic activity, stability and subcellular localization of the protein.

SIK2 has been implicated in the attenuation of insulin-mediated Akt signaling via phosphorylation of IRS1 on Ser789 in adipose tissue [30]. We previously showed that serine phosphorylation of IRS1 by SIK2 prevents insulin-induced Müller cell survival by downregulating the PI3K/Akt pathway [38]. It is possible that SIK2 participates in cross talk between the PI3K/Akt and the ERK pathways. SIK2 participation in a novel CREB-dependent survival response of cortical neurons has also been proposed [60]. Whether these mechanisms represent independent processes or are interconnected remains to be investigated.

Müller glia are structurally and functionally closely associated with retinal neurons, and they undergo dysregulation in the face of injury and disease states, a process known as reactive gliosis similar to astrogliosis seen elsewhere in the CNS [12]. Though this process is poorly understood, several growth factors including FGF2 and increased levels of ERK activity have been implicated in its development [12]. Reactive gliosis is thought to be an attempt to maintain tissue homeostasis [12]. Depending on the severity and the duration of the insult however changes in the metabolism, survival and proliferative state of these cells often exacerbate the pathology. Therefore, it will be of interest whether SIK2 plays similar roles in regulation of proliferation and survival of glia in vivo and if it contributes to reactive gliosis.

As Gab1 and IRS1 are structurally and functionally very similar proteins and partake in the signal transduction downstream of numerous RTK pathways, it is conceivable that SIK2 may be a common regulator of these pathways. Thus, perturbation of SIK2-dependent regulation of signaling pathways might have an impact on a multitude of biological processes. The negative regulatory role of SIK2 on proliferation observed here also raises the possibility that it acts as a tumor suppressor and that deficiencies in SIK signaling contribute to tumorigenesis. In line with this possibility an oncomine database search (www.oncomine.com) revealed reduced SIK2 levels in a number of cancer transcriptome studies. SIK2 was reported as a potential tumor suppressor in breast cancer *via* inhibiting PI3K/Akt and Ras/ERK signaling cascades simultaneously [75].

In contrast to tumor suppressor role, some reports suggesting an oncogenic role of SIK2 in ovarian [49,71], prostate [8], osteosarcoma [43] and colorectal [46] cancers via regulating different cellular mechanisms. The discrepancies between our results and these reports may reflect cellular and pathway specific aspects of SIK2 regulation.

## Conclusion

In summary, we have provided evidence for SIK2 involvement in FGF signal transduction and a new negative feedback loop that modulates FGF2-dependent ERK activation (Figure 7). We suggest that upon FGF2 stimulation activated ERK rapidly targets SIK2 on S/T residues within the putative phosphorylation motif (S/T)P and thereby enhances its catalytic activity. In turn, active SIK2 phosphorylates Gab1 on serine 266 residue, which decreases Grb2 and Shp2 association with Gab1. Loss of this interaction then leads to downregulation of ERK signaling, with the consequence of reduced proliferation.

**Figure 7.**
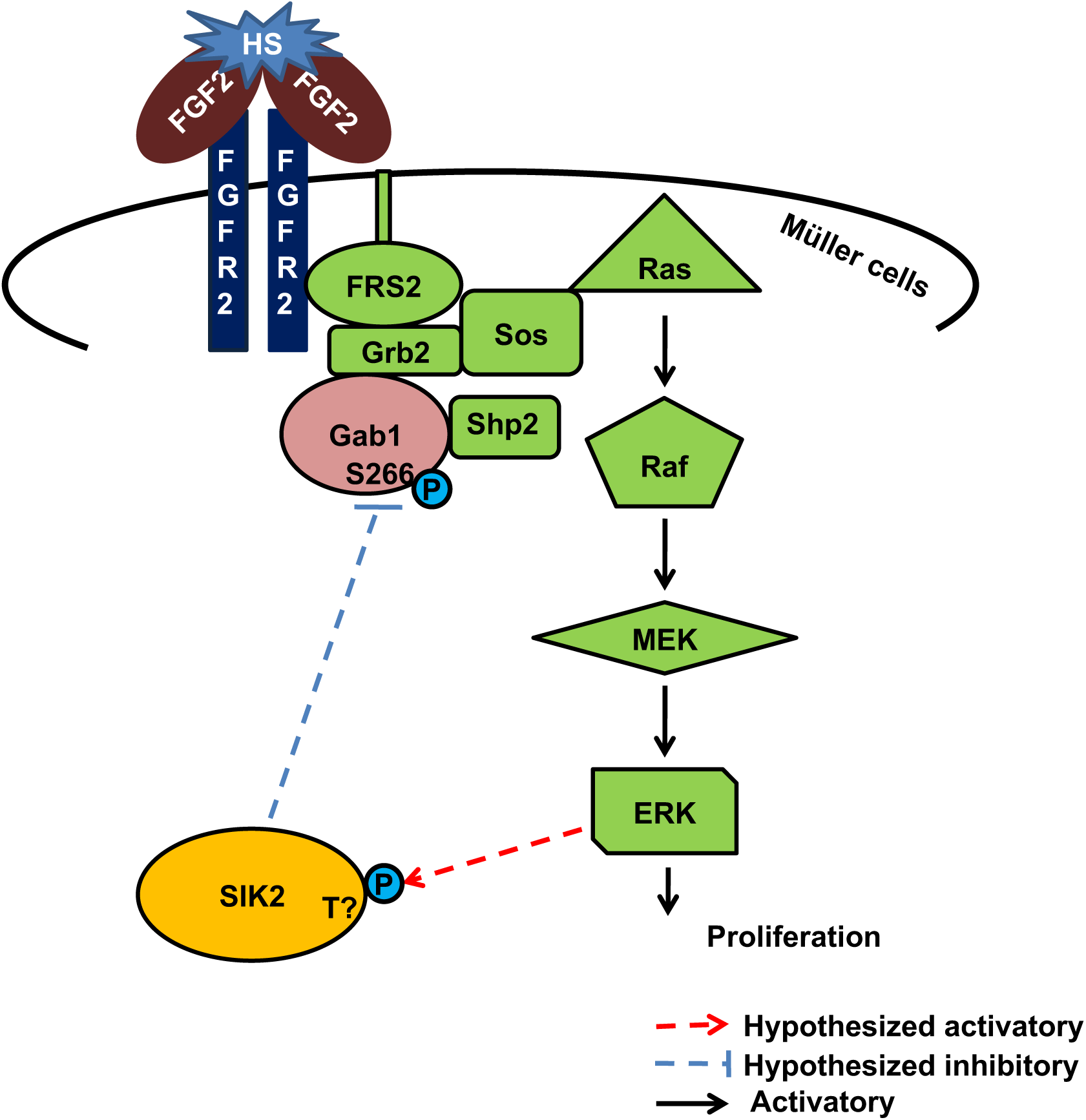
The proposed model. We propose that upon FGF2 stimulation activated ERK rapidly targets SIK2 on S/T residues and enhances its catalytic activity. In turn, active SIK2 phosphorylates Gab1 on serine 266 residue that hampers Grb2 and Shp2 association to Gab1. Thus, ERK signaling is downregulated in a feedback fashion resulting in reduced proliferation.

## Acknowledgements

Authors would like to thank to Dr. David Hicks (Université de Strasbourg, France), Dr. Stefan Fuss and Dr. İbrahim Yaman (Bogazici University, Turkey) for critically reading and valuable comments during preparation of the manuscript. This work was supported by grants from the Bogazici University Research Projects (5581); Turkish Scientific and Research Council (109T200; 106T115). Authors acknowledge the use of Biorender to generate graphical abstract.

